# Pancreatic Stellate Cells Secrete Deoxycytidine Conferring Resistance to Gemcitabine in PDAC

**DOI:** 10.1101/558627

**Authors:** Simona Dalin, Mark R. Sullivan, Emanuel Kreidl, Allison N. Lau, Beatrice Grauman-Boss, Silvia Fenoglio, Alba Luengo, Matthew G. Vander Heiden, Douglas A. Lauffenburger, Michael T. Hemann

## Abstract

Pancreatic ductal adenocarcinoma (PDAC) is a leading cause of cancer deaths in the United States. The deoxynucleoside analog gemcitabine is among the most effective therapies to treat PDAC; however, nearly all patients treated with gemcitabine either fail to respond or rapidly develop resistance. One hallmark of PDAC is a striking accumulation of stromal tissue surrounding the tumor, and this accumulation of stroma can contribute to therapy resistance. To better understand how stroma limits response to therapy, we investigated cell-extrinsic mechanisms of resistance to gemcitabine. We show that conditioned media from pancreatic stellate cells (PSC), as well as from other fibroblasts, protects PDAC cells from gemcitabine toxicity. We find that the PSC conditioned media protective effect is mediated by secretion of deoxycytidine, but not other deoxynucleosides, through equilibrative nucleoside transporters. Deoxycytidine inhibits the processing of gemcitabine in PDAC cells, thus reducing the effect of gemcitabine and other nucleoside analogs on cancer cells. Our results suggest that reducing deoxycytidine production in PSCs may increase the efficacy of nucleoside analog therapies.

**Additional Information:** Funding: This project was funded in part by the NIH (NCI U54-217377), the MIT Center for Precision Cancer Medicine, and by the Koch Institute Support (core) Grant P30-CA14051 from the National Cancer Institute. S.D. was supported by the David H. Koch Fellowship in Cancer Research. A.N.L was a Robert Black Fellow of the Damon Runyon Cancer Research Foundation, DRG-2241-15 and was supported by a NIH Pathway to Independence Award (K99/R00), 1K99CA234221. M.T.H and M.G.V.H. acknowledges funding from the MIT Center for Precision Cancer Medicine and the Ludwig Center at MIT. M.G.V.H also acknowledges funding from the Lustgarten Foundation, SU2C, the MIT Center for Precision Cancer Medicine, the NCI, and an HHMI Faculty Scholar award.

Competing interests: M.G.V.H. is a consultant and advisory board member for Agios Pharmaceuticals, Aeglea Biotherapeutics, and Auron Therapeutics.

## Introduction

Pancreatic ductal adenocarcinoma (PDAC) is notorious as a cancer with poor therapeutic options for most patients. Few patients with PDAC are eligible for surgery, and systemic therapies are all limited in their benefit. FOLFIRINOX combination therapy (Folinic acid, 5-fluorouracil, irinotecan and oxaliplatin) is the most effective regimen for patients who can tolerate the significant side effects. Gemcitabine also can be effective in some patients and has more tolerable side effects *(1)*. Regardless of which therapy is used, the 5-year survival rate is only 8.5%, and, due to increasing incidence, PDAC is projected to become the second leading cause of cancer deaths by 2030 *(2, 3)*.

One striking feature of PDAC is a significant accumulation of stiff stromal tissue surrounding the tumor, termed the desmoplastic response, and this has been proposed as one factor that makes this cancer so resistant to therapy *(4–6).* Initially, the stiff stroma was thought to present a physical barrier to drug perfusion, however it is now appreciated that limiting drug delivery is not the only reason pancreatic tumors are therapy resistant *(6–8)*. The ways in which the stromal cells within pancreatic cancer contribute to PDAC resistance has become an area of intense study, with a recent focus on interaction between stroma and PDAC via direct contact and paracrine signaling *(9–12).* A variety of biomolecules secreted by pancreatic stromal cells have been implicated in resistance to radio- and chemotherapy, including cytokines such as SDF-1a and IGFs, extracellular matrix (ECM) proteins such as fibronectin and collagen I, and even exosomes *(13–17)*.

In addition to the above methods, the stroma has also been shown to cause resistance by sequestering gemcitabine, a deoxycytidine (dC) analog *(11, 18)*. The mechanism by which gemcitabine causes toxicity in cells in well studied *(19)*. This drug enters cells through a variety of nucleoside transporters *(20)*. Once inside cells, gemcitabine is activated via phosphorylation by deoxycytidine kinase to form the active gemcitabine-triphosphate, which can be incorporated into DNA in place of dCTP resulting in masked chain termination, DNA damage, and apoptosis *(21)*.

Here, we show that PSCs, a pancreatic stromal cell type, secrete dC through equilibrative nucleoside transporters. In turn, deoxycytidine protects PDACs from gemcitabine toxicity. We show that dC-mediated gemcitabine resistance likely acts at the level of competition for deoxycytidine kinase. Several other fibroblast cell lines also secrete dC, and media conditioned by PSCs cause resistance to other dC analogs as well as deoxyadenosine analogs. Targeting this stromal dC production and secretion may represent an opportunity to improve the efficacy of nucleoside analog therapies.

## Materials and Methods

### Cell Culture

P53 2.1.1 PDAC cells derived from *p48^cre^, Kras^LSL_G12D/wt^, Tp53^flox/wt^* mice were obtained from Eric Collison (UCSF) *(22)*. PSC, PSC1, and PSC3 cells were derived from the pancreas of FVB/nj mice upon induction of pancreatitis via caerulein treatment mice as described *(23, 24)*. In brief, mice were injected with 8 hourly injections of 50μg/kg of caerulein for two consecutive days. Mice were housed animal research facilities of the MIT Division of Comparative Medicine (DCM). All facilities are fully accredited by the AAALAC (Animal Welfare Assurance number A-3125) and meet NIH standards as set forth in the “Guide for Care and Use of Laboratory Animals” (DHHS). The MIT DCM Committee on Animal care last reviewed and approved the planned research in this proposal for the Hemann lab (Protocol Number 0515-044-18) on 4/12/18. After the last injection, pancreata were dissected and minced into 2-3 mm pieces which were cultured in a 10 cm dish in 10 mL of DMEM:F12 medium supplemented with 2 mM glutamine and 15% fetal bovine serum. After 72 hours, tissue pieces were removed and the PSCs were immortalized by transduction with SV40. The infected population was selected by puromycin treatment at 2 μg/mL. PSC6 cells were derived from the pancreas of C57BL/6J mice via density centrifugation as previously described *(25)*, and were immortalized as described above. 293T cells were purchased from ATCC. Hepatic stellate cells (HSC) were a gift from Dr. Linda Griffith (MIT). Primary and immortalized MEFs were isolated and cultured as previously described *(24)*. Cell lines were maintained in DMEM (Corning) supplemented with Penicillin-Streptomycin (Corning), 2mM L-glutamine (Corning), and 10% FBS (HyClone) and routinely tested for mycoplasma contamination using MycoAlert (Lonza).

### Conditioned Media

Conditioned media (CM) was produced by changing culturing media of 70%-90% confluent plates, then collecting the media after the indicated number of days passed. CM was filtered through a 0.2 μM SFCA membrane (Corning). Unless otherwise noted, cells conditioned the media for three days prior to collection, and CM was used at a 1:5 dilution. When not in use, CM was stored at −20°C. Boiled CM was incubated at 100°C for 1 hour and centrifuged to remove precipitate. Proteinase K digested CM was incubated with 2 g/mL proteinase K (Invitrogen) for 1 hour at 37°C. Proteinase K was inactivated at 95°C for 10 minutes and the digested media was centrifuged to remove precipitate. Size cutoff columns (EMD Millipore) were used to filter out all compounds larger than 3kDa from the media.

### Chemicals

Gemcitabine, azacytidine, and decitabine were purchased from LC Labs. Fludarabine, azidothymidine, and cladribine were purchased from Tocris. Cytarabine was purchased from Selleck Chemicals. Puromycin, nucleosides, nucleotides, and dCTP isotope were purchased from Sigma. Isotopes of deoxycytidine, deoxythymidine, deoxyguanosine, and deoxyadensoine were purchased from Cambridge Isotope Laboratories. Deoxyuridine isotope was purchased from Santa Cruz. Cytarabine, decitabine, azidothymidine, and azacytidine were dissolved in DMSO (Sigma) at 50-100mM and stored at −80°C until use. Working stocks were diluted to 5mM in 100% ethanol (VWR) and stored at −20°C until use. Fludarabine and Cladribine were dissolved in DMSO at 50mM, then divided to 100 μL aliquots. Aliquots were stored at −80°C. Aliquots were diluted in DMSO to 5mM and divided to 10 μL aliquots and stored at −20°C, to reduce freeze/thaw cycles. Puromycin was dissolved in sterile water at 50mM and stored at −20°C. Gemcitabine was dissolved in DMSO at 10mM and stored at −20°C. Nucleosides were dissolved in sterile water at 10mM and stored at −80°C. Nucleoside and nucleotide isotopes were dissolved in HPLC-grade water (Sigma) at 1mM and stored at −80°C.

### Cell Viability Assay

Cells were seeded in cell-culture treated 384-well plates (Falcon) in triplicate and spun for 5 minutes at 1500 RPM then allowed to attach for 4-6 hours. After attachment, cells were treated as indicated. Cell viability was measured after 72 hours using resazurin sodium salt (Sigma). Resazurin was used at 0.167 mg/mL and plates were analyzed 3-4 hours after addition. Fluorescence was measured using the Tecan M200 Pro at an excitation of 550nm and emission of 600nm. GR50 values were calculated using GraphPad Prism 5.

### Metabolite Extraction

For analysis of conditioned media or adherent cells, cells or 10 μL of media were combined with 600 μL HPLC grade methanol (Sigma-Aldrich, 646377-4X4L) containing 1 μM each of ^15^N3 ^13^C9-dCTP, ^13^C1 ^15^N2-deoxyuridine, ^13^N3-deoxycytidine, ^13^C10 ^15^N2-thymidine, ^15^N5-deoxyguanosine, and ^15^N5-deoxyadenosine, 300 μL HPLC grade water (Sigma-Aldrich), and 400 μL HPLC grade chloroform (Sigma-Aldrich). Samples were vortexed for 10 minutes at 4 °C, then centrifuged at 21000 x g at 4°C for 10 minutes. 400 μL of the aqueous layer was removed and dried under nitrogen.

### HPLC fractionation

Conditioned media was extracted as described above, then the dried, aqueous layer was resuspended in 65 μL 80:20 acetonitrile:water. 40 μL of the resuspended liquid was then fractionated using an Agilent 1200 Series HPLC. Fractionation was done using a Luna 5 μm, 100 x 4.6 mm HILIC column with 200 Å pore size (Phenomenex 00D-4450-E0). Mobile phase A was 100 mM ammonium formate, pH 3.2. Mobile phase B was 100% HPLC grade acetonitrile (Sigma-Aldrich). The chromatographic gradient was: 0-3.5 minutes: hold at 90% mobile phase B; 3.5-11 minutes: linear gradient from 90% to 50% mobile phase B; 11-13.5 minutes: hold at 50% mobile phase B; 13.5-16 minutes: linear gradient from 50% mobile phase B to 90% mobile phase B. Flow rate was 1 mL/minute. Fractions were collected in increments of one minute. Each fraction was lyophilized with a Labconco FreeZone 2.5, then resuspended in 40 μL of 44 mM sodium bicarbonate at pH 7.4. Fractions were used directly in LC/MS experiments as described below, or supplemented with 10% FBS for use in cell viability assays with gemcitabine.

### LC/MS

Dried media and cell extracts were resuspended in 100 μL HPLC grade water. LC/MS analysis was performed using a QExactive orbitrap mass spectrometer using an Ion Max source and heated electro-spray ionization (HESI) probe coupled to a Dionex Ultimate 3000 UPLC system (Thermofisher). External mass calibration was performed every 7 days. Samples were separated by chromatography by injecting 10 μL of sample on a SeQuant ZIC-pHILIC 2.1 mm x 150 mm (5 μm particle size) column. Flow rate was set to 150 mL/min. and temperatures were set to 25°C for the column compartment and 4°C for the autosampler tray. Mobile phase A was 20 mM ammonium carbonate, 0.1% ammonium hydroxide. Mobile phase B was 100% acetonitrile. The chromatographic gradient was: 0–20 min.: linear gradient from 80% to 20% mobile phase B; 20–20.5 min.: linear gradient from 20% to 80% mobile phase B; 20.5 to 28 min.: hold at 80% mobile phase B. The mass spectrometer was operated in full scan, polarity-switching mode and the spray voltage was set to 3.0 kV, the heated capillary held at 275°C, and the HESI probe was held at 350°C. The sheath gas flow rate was 40 units, the auxiliary gas flow was 15 units and the sweep gas flow was one unit. The MS data acquisition was performed in a range of 70–1000 m/z, with the resolution set at 70,000, the AGC target at 1×10^6^, and the maximum injection time at 20 msec. Relative quantitation of polar metabolites was performed with XCalibur QuanBrowser 2.2 (Thermo Fisher Scientific) using a 5 ppm mass tolerance and referencing an in-house library of chemical standards. For analysis of deoxynucleoside and dCTP concentrations in conditioned media, peak areas were compared to stable isotope labeled internal standard peak areas to calculate concentration. For analysis of intracellular metabolite concentrations, peak areas were normalized using ^13^C stable isotope labeled internal standards and intracellular concentrations were calculated using measurements of total cell volume made using a Coulter counter.

### Statistical Analysis

Statistics were performed using GraphPad Prism 5 (GraphPad Software Inc). All error bars represent standard error of the mean.

## Results

### PSCs secrete a small molecule that protects PDAC cells from gemcitabine toxicity

To better understand tumor microenvironmental contributions to gemcitabine resistance in PDAC, we extracted activated PSCs via the outgrowth method as previously described *(23, 24).* We conditioned media (CM) in the presence of these PSCs for varying times and then examined gemcitabine dose responses on PDAC cells in the presence of varying dilutions of this CM (Fig. 1A). Media conditioned by the PSCs for 3 days conferred 11-fold increase in the dose of gemcitabine required to achieve 50% reduction in the growth rate of PDAC cells (GR50, a metric which takes into account cytostatic and cytotoxic effects of a pertubation) *(25)*. Further, this protective effect was present even in highly diluted CM, with 5% CM conferring 2-fold increase in GR50 (Fig. 1B). The protective effect accumulated in the CM over time, conferring almost 100-fold increase in gemcitabine GR50 after 9 days of media conditioning (Fig. 1C). The CM also did not affect the proliferation of PDAC cells in the absence of gemcitabine. To begin to characterize the activity in conditioned media that is responsible for this effect, we incubated the CM at 100°C for 1 hour, digested the CM with proteinase K, and tested the flow through of CM passed through a 3 kDa cutoff filter. The protective effect of the CM was retained even after boiling, proteinase K exposure, or passage through a filter to deplete high molecular weight material, implicating a small metabolite as the likely cause of the protective effect (Fig. 1D).

**Fig. 1.**
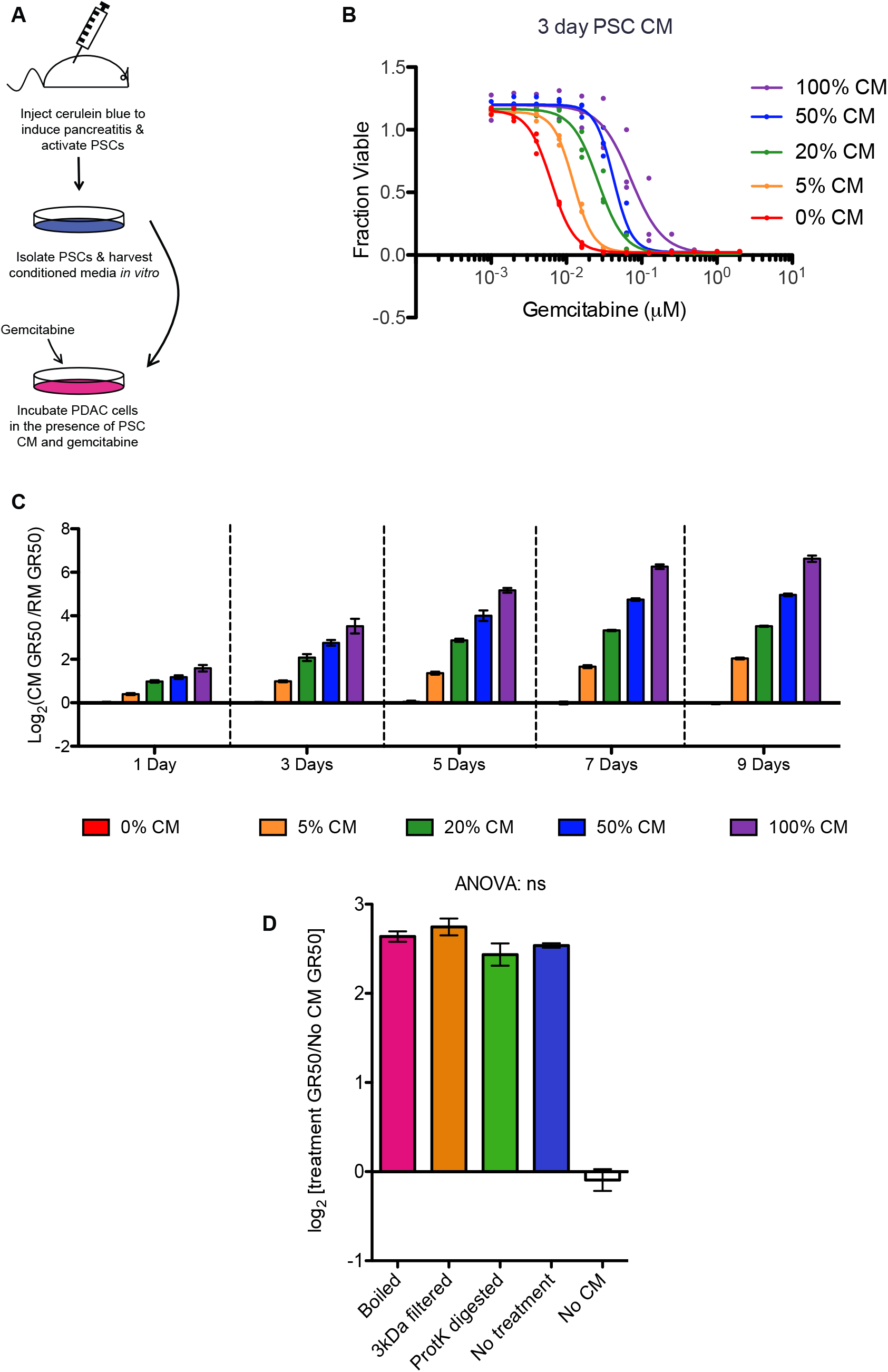
PSCs secrete a small metabolite that causes PDAC resistance to gemcitabine. **(A)** Schematic of generating conditioned media and testing the protective effect. **(B)** Gemcitabine dose response curves on PDAC cells supplemented with the indicated percentage of PSC CM harvested after 3 days in culture. Three biological replicates are shown. **(C)** Log2 of the gemcitabine GR50 fold change induced by supplementing the indicated percentage of PSC CM produced for the indicated number of days. Data represent the mean of three biological replicates ± SEM. **(D)** Log2 fold changes of gemcitabine GR50s on PDAC cells supplemented with 3-day PSC CM treated as indicated. Three biological replicates are shown. Data represent the mean of three biological replicates ± SEM. ns – not significant (one-way ANOVA).

### The protective small metabolite can be fractionated by HPLC

To identify the small metabolite secreted by PSCs that causes resistance to gemcitabine, we fractionated PSC CM using HILIC HPLC chromatography. After buffer removal via lyophilization, we diluted each fraction 1:5 and tested the protective effect of the CM on cells treated with 0.02 μM gemcitabine. After three days at this dose, there are about half the number of live cells in the absence of conditioned media relative to untreated controls. In the presence of PSC CM diluted to match the amount found in each HPLC fraction, there are ~85% of the number of viable cells compared to untreated controls (Fig. 2A). The number of viable cells in fraction 4 of conditioned media was 79.1% relative to untreated controls, indicating the presence of the protective activity in fraction 4. To confirm that the fractionation isolated activity that was not resulting simply from byproducts of cell growth, we used the same protocol to fractionate 293T CM, which does not have any protective effect. Fractionated 293T CM had 60% or fewer viable cells compared to untreated controls in all fractions (Fig. 2A).

**Fig. 2.**
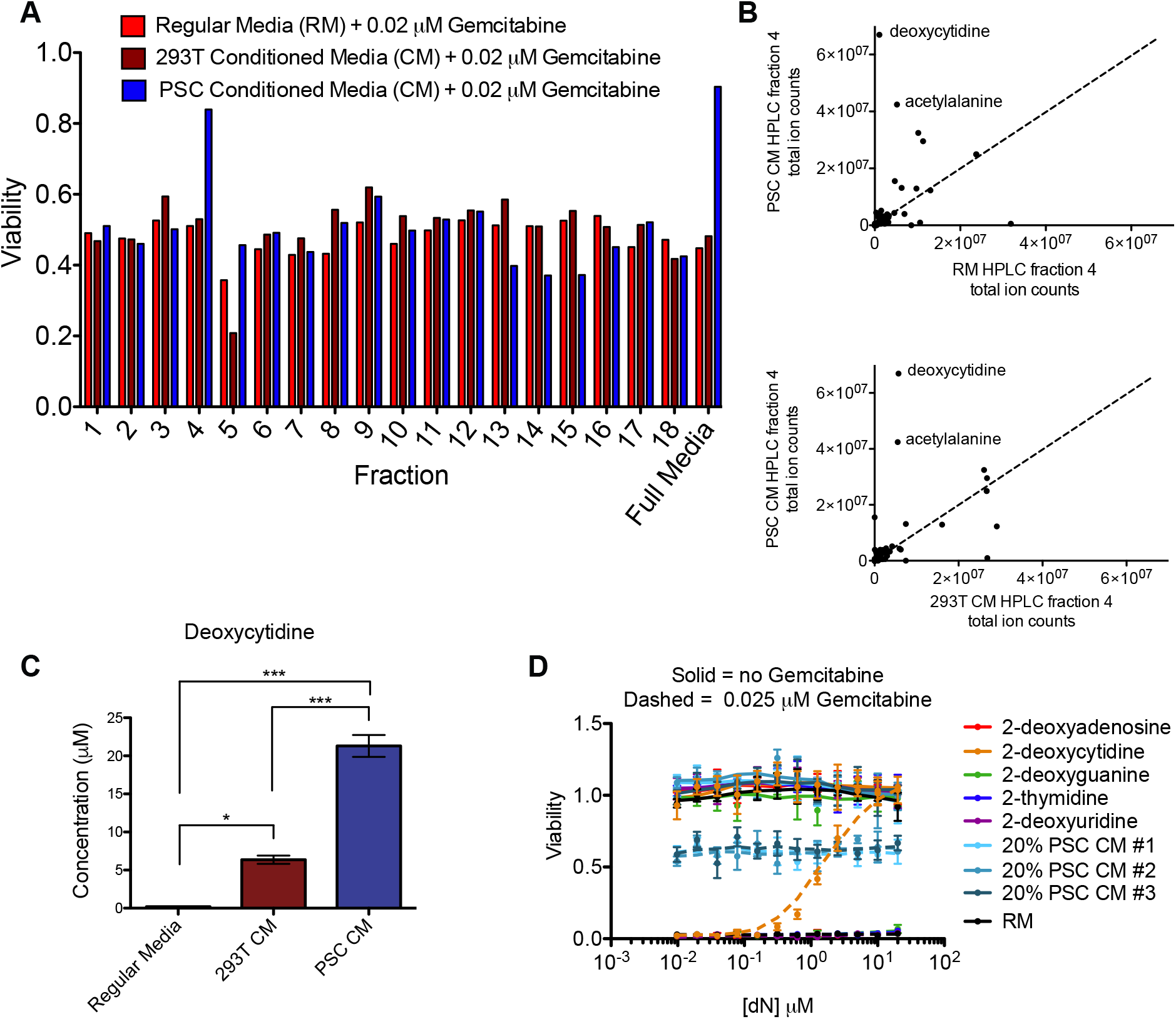
Deoxycytidine is present in PSC CM and is sufficient to protect PDAC cells from gemcitabine toxicity. **(A)** Viability of PDAC cells treated with 0.02 μM gemcitabine and the indicated HPLC fraction of regular media (RM), 3-day 293T CM, or 3-day PSC CM. **(B)** A scatterplot of ion counts of 150 metabolites in fraction 4 of RM vs. 3-day PSC CM (top) and 3-day 293T CM vs. 3-day PSC CM (bottom). **(C)** Quantification of deoxycytidine in RM, 3 day 293T CM, and 3-day PSC CM. Data represent the mean of three biological replicates ± SEM. *, P ≤ 0.05, ***, P ≤ 0.001 (one-way ANOVA with post hoc Bonferroni-tests) **(D)** Dose response curves with the indicated nucleosides without (solid lines) or with (dashed lines) 0.025 μM gemcitabine. The viability of 20% 3-day PSC CM is also shown. Data represent the mean of three biological replicates ± SEM.

### Deoxycytidine accumulates in PSC conditioned media

To identify the activity responsible for the protective effect observed in HPLC fraction 4, we used liquid chromatography/mass spectrometry (LC/MS) to assess the abundance of 150 common plasma metabolites present in HPLC fraction 4 from PSC CM, 293T CM, and regular non-conditioned media (RM). Of these metabolites, deoxycytidine (dC) and acetylalanine were the only compounds present in higher quantities in fraction 4 of PSC CM compared to the equivalent fraction from 293T CM and RM (Fig. 2B). N-acetyl alanine was unable to protect PDAC cells from gemcitabine toxicity at doses up to 50 μM (Fig. S1A), so we focused our analysis on dC. Interestingly, after three days in culture, dC in PSC CM accumulated to 21.3 μM compared to 0.18 μM in RM and 6.4 μM in 293T CM (Fig. 2C), suggesting that PSCs may be exporting deoxycytidine into the media.

### Deoxycytidine is sufficient to protect PDAC cells from gemcitabine toxicity

To determine if dC or other nucleosides are sufficient to protect PDAC cells from gemcitabine toxicity, we conducted dose response curves with each nucleoside in the presence or absence of 0. 025 μM gemcitabine, a dose that kills 100% of PDAC cells after 3 days in the absence of CM. Of all five nucleosides tested, only dC protected PDAC cells from gemcitabine toxicity, with a GR50 of 0.46 μM confirming the sufficiency of dC to cause the protective phenotype (Fig. 2D).

### PSCs do not produce substantial quantities of other nucleosides

While only dC protected PDAC cells from gemcitabine toxicity, we asked if PSCs secrete other nucleosides as well. Using LC/MS, levels of thymidine, deoxyuridine, dC, deoxyguanosine, and dCTP were quantitated in 3-day PSC CM as well as in metabolite extracts of PSC cells. We confirmed high levels of dC secretion into the media, and also observed high levels of intracellular dC at 88 μM. PSCs contained detectable levels of thymidine and deoxyuridine with some release into the media albeit at lower levels than dC. Deoxyguanosine (dG) was detected in cells, but not in the media. dCTP was also only detected in cells and not in the media (Fig. 3A,B).

**Fig. 3.**
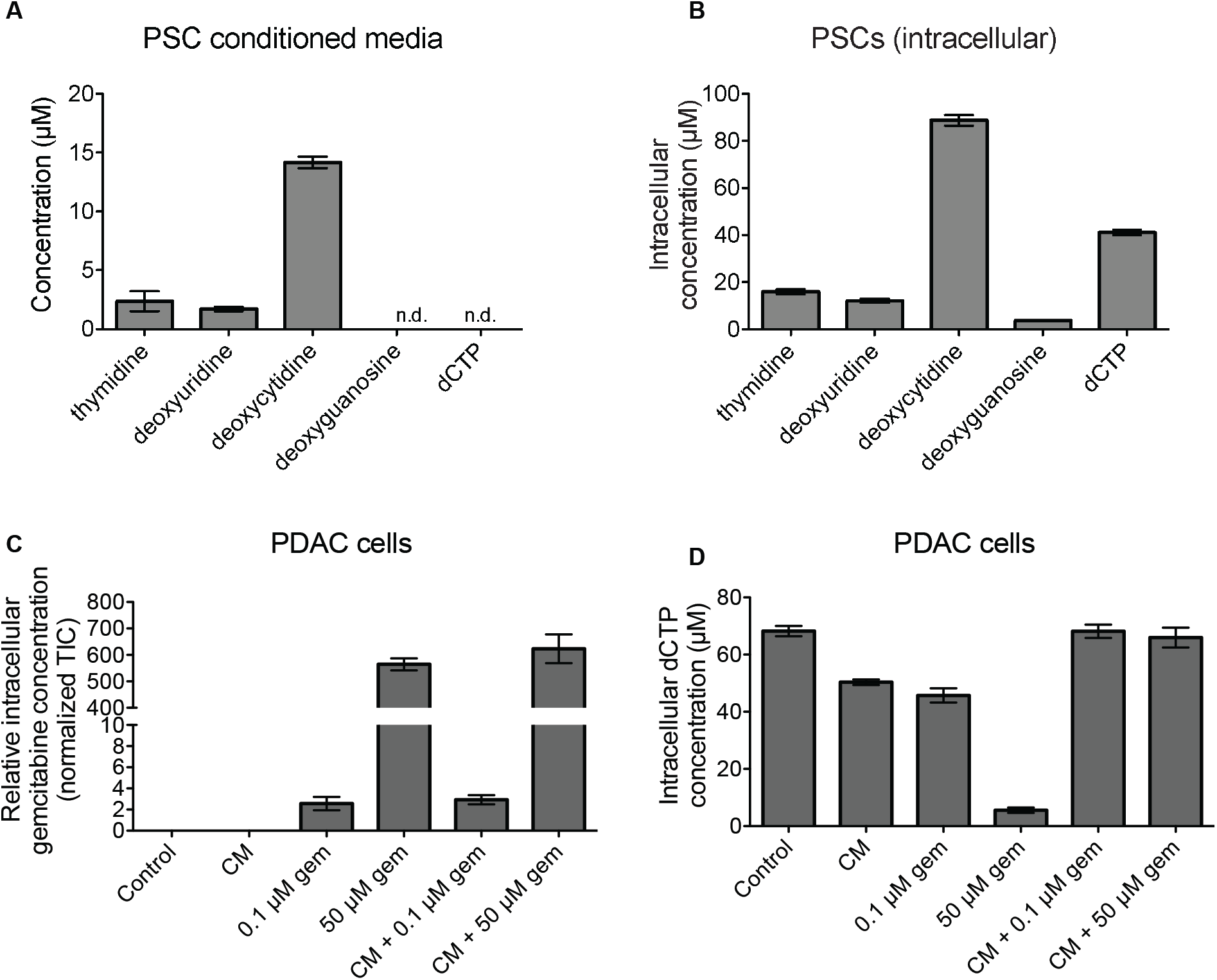
PSCs produce and secrete deoxycytidine at higher levels than other nucleosides. **(A)** Absolute quantification of the indicated species in 3-day PSC CM. Data represent the mean of 3 biological replicates ± SEM. Undetected species are indicated by “n.d.” **(B)** Absolute quantification of the indicated species in PSC extracts after producing CM for 3 days. Data represent the mean of 3 biological replicates ± SEM. **(C)** Relative quantification of gemcitabine within PDAC cells after the indicated treatment. Data represent the mean of 3 biological replicates ± SEM. **(D)** Absolute quantification of dCTP within PDAC cells after the indicated treatment. Data represent the mean of 3 biological replicates ± SEM.

### PSC conditioned media does not prevent gemcitabine from entering PDAC cells

Gemcitabine is transported into cells via the same nucleoside transporters that transport dC *(21)*. Thus, we hypothesized that the dC present in the PSC CM may compete with gemcitabine for these transporters, blocking gemcitabine from entering the cell. We treated PDAC cells with 0.1 μM gemcitabine, 50 μM gemcitabine (representative of patient plasma concentration achieved after infusion *(26)*), and/or diluted PSC CM for two hours. We then quantified intracellular gemcitabine levels, as well as thymidine, deoxyuridine, dC, deoxyguanosine, and dCTP using LC/MS. Interestingly, treatment with PSC CM did not affect intracellular gemcitabine levels at either dose tested, suggesting that deoxycytidine competes with gemcitabine downstream of uptake via nucleoside transporters (Fig. 3C). Similarly, gemcitabine treatment did not affect import of dC or other nucleosides from PSC CM into PDAC cells (Fig. S1B-E). Treatment with PSC CM did rescue intracellular dCTP levels after treatment with 50 μM gemcitabine, suggesting that PSC CM is able to protect cells from gemcitabine toxicity even at this high physiologic dose (Fig. 3D).

### Inhibiting equilibrative nucleoside transporters on PSCs reduces deoxycytidine in conditioned media

We hypothesized that the PSCs were secreting large quantities of dC through equilibrative nucleoside transporters (ENTs) which have previously been shown to transport dC *(20)*. Indeed, when we treated PSCs with a non-toxic dose of two different ENT inhibitors (Fig. S2A), the final concentration of deoxycytidine in the CM from these cells was reduced to a level similar to that in 293T CM (Fig. 4A, Fig. S2B). Treatment with ENT inhibitors did not effect levels of other nucleosides secreted by PSCs (Fig. S2 C-D). Gemcitabine can be transported into PDAC cells through the same transporters and we were unable to inactivate ENT inhibitors by boiling the CM (Fig. S3), likely explaining why ENT inhibitor treatment did not completely abolish the ability of PSC CM to protect cancer cells from gemcitabine toxicity (Fig. 4B).

**Fig. 4.**
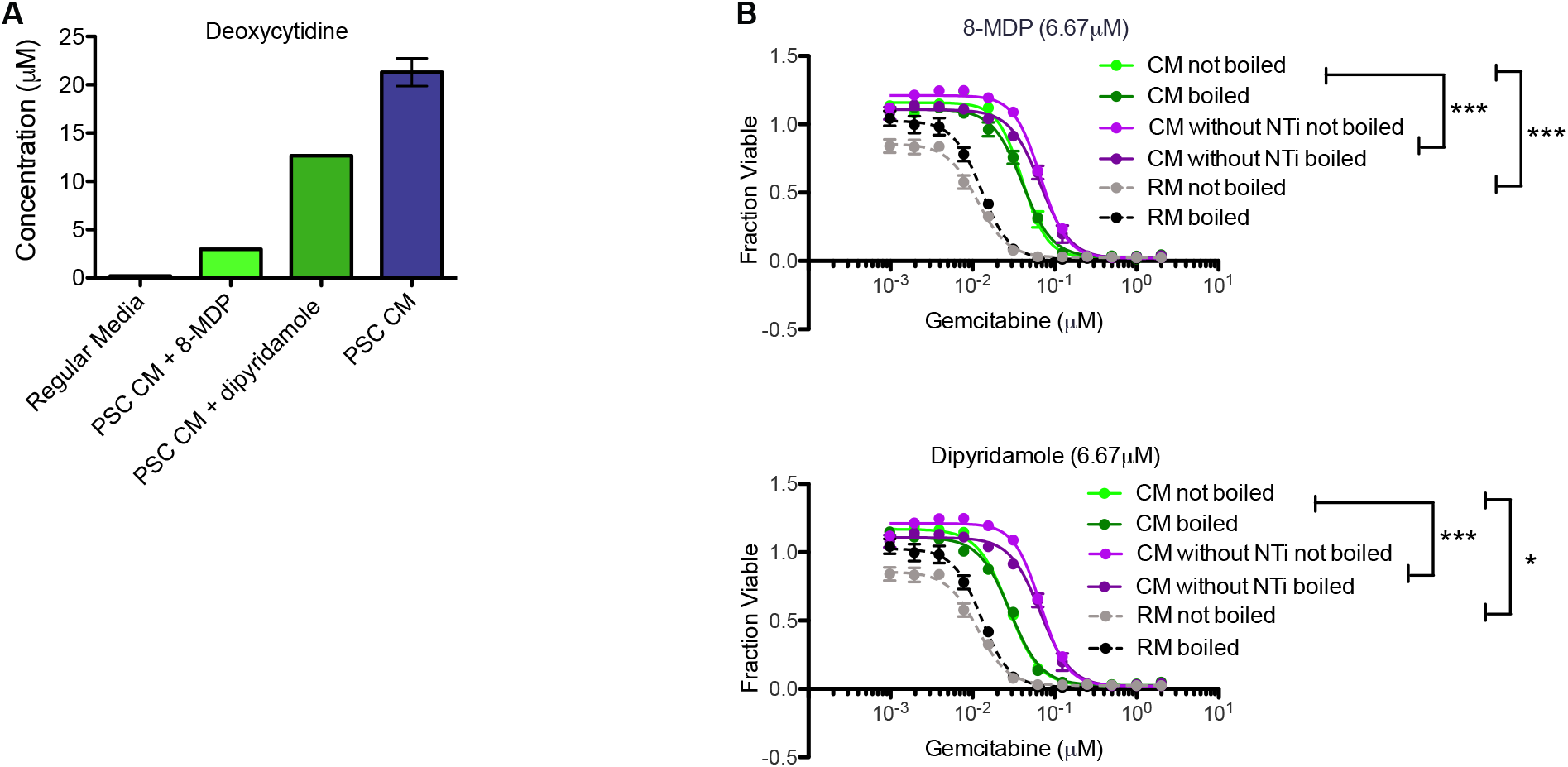
PSCs secrete dC through ENTs. **(A)** The deoxycytidine concentration in 3-day CM from PSCs treated with 6.67 μM of the indicated ENT inhibitors is shown. Data represent the mean of three biological replicates ± SEM except for 8-MDP and dipyridamole treated CM which represent one biological replicate. **(B)** Gemcitabine dose response curves of PDAC cells supplemented with PSC conditioned media made in the presence of the indicated ENT inhibitor. Data represent the mean of three biological replicates ± SEM. *, P ≤ 0.05, ***, P ≤ 0.001 (one-way ANOVA with Bonferroni post-tests).

### Some other fibroblast cell types also secrete deoxycytidine

In order to determine if this protective effect is specific to PSCs, we analyzed CM from several additional PSC cell lines isolated with different methodologies or from different mouse background (PSC1, PSC3, PSC6), mouse embryonic fibroblasts (MEFs), and hepatic stellate cells (HSCs). We found that CM from the additional PSC cell lines and MEFs did protect PDAC cells from gemcitabine toxicity to varying degrees. However, HSC CM did not protect PDACs from gemcitabine toxicity (Fig. 5A), suggesting secretion of deoxycytidine is not unique to PSCs, but not all fibroblast cell types secrete deoxycytidine.

**Fig. 5.**
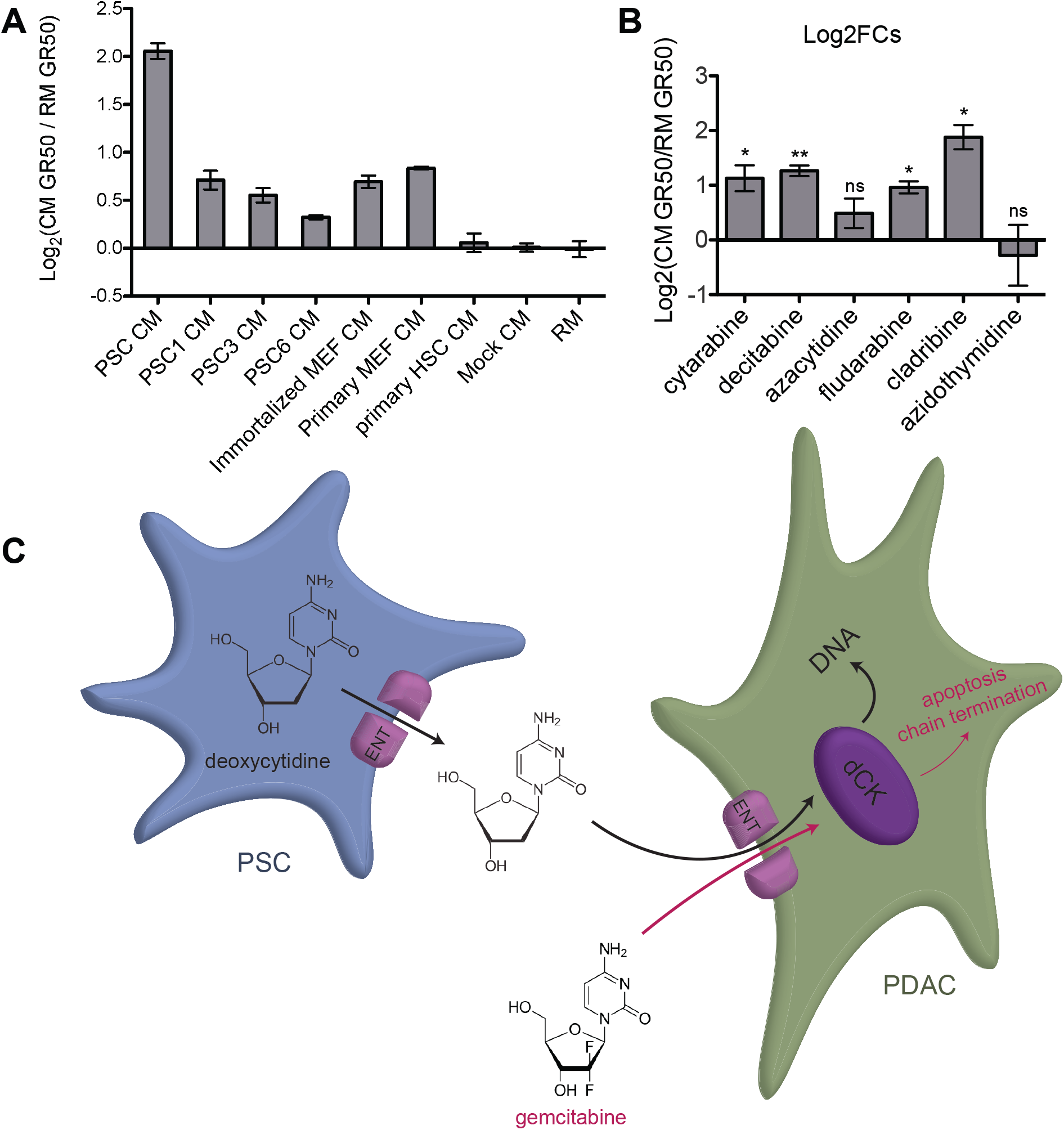
Other PSCs and MEFs protect PDACs from gemcitabine toxicity, and CM is active against other nucleoside analogs. **(A)** The log2 fold change in gemcitabine GR50 after treatment with 3-day CM from the indicated cell line is plotted. Data represent the mean of three biological replicates ± SEM. **(B)** Log2 fold changes of GR50s of the indicated nucleoside analog drugs supplemented with 3-day PSC. Data represent the mean of three biological replicates ± SEM. *, P ≤ 0.05, **, P ≤ 0.01 (two-tailed one sample t-test). **(C)** Diagram of proposed model. PSCs produce and secrete deoxycytidine through ENTs. Gemcitabine and deoxycytidine are transported into PDACs through ENTs and compete for phosphorylation by dCK.

### PSC conditioned media protects PDACs from other nucleoside analog drug toxicity

Other nucleoside analogs are active chemotherapy agents, and we conducted dose response curves to determine if the PSC CM can protect against other nucleoside analog toxicity. As expected, diluted PSC CM provided 2-fold increase in GR50 to the deoxycytidine analogs cytarabine and decitabine. Interestingly, the CM also caused a 2-fold increase in GR50 to deoxyadenosine analogs fludarabine and cladribine. Toxicity of the cytidine analog azacytidine and the thymidine analog azidothymidine were not affected by the CM (Fig. 5B). These data suggest that production of dC by resident fibroblasts might contribute to cancer resistance to some nucleoside analog chemotherapies.

## Discussion

Our results add deoxycytidine to the arsenal of molecules secreted by pancreatic stroma that contribute to chemotherapy resistance in PDAC. Deoxycytidine does not hinder gemcitabine entry into cells, suggesting competition for downstream processing enzymes such as deoxycytidine kinase. Deoxycytidine secretion may be a conserved phenotype of some types of fibroblasts, as MEF CM was able to protect PDAC cells from gemcitabine toxicity, albeit to a lesser extent. However, the lack of dC secretion by hepatic stellate cells suggests this is not a property of all fibroblasts and other factors might influence whether these support cells produce this molecule.

While PSC CM did not protect PDAC cells from azacytidine or azidothymidine toxicity, it did confer protection against cytarabine, decitabine, fludarabine, and cladribine. Neither azacytidine nor azidothymidine rely on deoxycytidine kinase for processing into their toxic forms, but cytarabine, decitabine, fludarabine, cladribine, and gemcitabine are all phosphorylated by deoxycytidine kinase into their toxic forms *(27, 28)*. This distinction further supports the hypothesis that nucleoside analog toxicity inhibition by deoxycytidine acts at the level of deoxycytidine kinase (Fig. 5C).

We observed that the high levels of PSC nucleoside production and secretion were limited to specifically deoxycytidine. Although we were unable to measure deoxyadenosine levels, deoxyguanosine was at a low concentration in PSCs, and was undetectable in PSC CM. This potential pyrimidine specificity suggests that the mechanism of deoxycytidine production is not due to general upregulation of nucleoside production. Deoxycytidine secretion is a known mechanism of maintaining intracellular dCTP pool sizes when too much is being produced *(29–31)*. Therefore, the deoxycytidine production and secretion specificity also suggests that PSCs may have increased levels of dCTP production perhaps as a biproduct of another metabolic process, and excess dCTP is converted into deoxycytidine and secreted.

The deoxycytidine specificity further suggests that the PDAC microenvironment may have a high level of deoxycytidine. At first glance this seems paradoxical – the microenvironment of PDAC is specifically protective against one of the only drugs effective for this tumor type, and which is one of only a small number of malignancies in which this drug is used *(1, 26)*. We propose instead that the behavior and microenvironment of tumors in a tissue dictates the drugs that are effective for that tumor type. We have previously observed this when comparing the variable efficacy of two platinum-based chemotherapies, cisplatin and oxaliplatin, in different tumor types. Cisplatin, which kills cells by inducing DNA damage, is more effective in breast and non-small cell lung cancers, where inactivating mutations in ATM and CHK2 predispose cells to DNA damage. On the other hand, oxaliplatin, which kills cells by inducing ribosome biogenesis stress, is more effective in gastrointestinal tumors which have increased expression of translation machinery *(32)*. In the pancreatic setting, PDAC cells may have a high need for deoxycytidine and therefore are optimized to import and utilize deoxycytidine and its analogs – explaining its efficacy in this tumor type. This may even suggest a high need for deoxycytidine in normal pancreatic ductal cell biology.

The high levels of deoxycytidine secreted by PSCs raises the question of *why* the PSCs are producing and secreting large quantities of this nucleoside. One hypothesis is that PSCs produce deoxycytidine as part of wound healing response, potentially in response to PDAC derived factors produced as a result of stress *(33–35)*. Regardless, these data motivate further understanding of nucleoside metabolism in fibroblasts and what drives dC secretion. Better understanding the mechanism of deoxycytidine production in PSCs could identify a way to increase the efficacy of nucleoside analog therapies in multiple cancer types.

## Acknowledgments

We thank Linda Griffith, for donation of the HSC cells. We are grateful to Chris Nabel and Jackie Lees for helpful discussion and to Sonya Entova for assistance with HPLC buffer removal.

## Author contributions

S.D., M.R.S., E.K., B.G.B, S.F., A.N.L., and S.E. performed the experiments. S.D., M.R.S., and E.K. analyzed the data. S.D. wrote the manuscript. S.D., M.R.S., E.K., M.G.V.H., D.A.L., and M.T.H conceived the project and designed the experimenthree biological replicatests.

## References and Notes

1. J. Kleeff, M. Korc, M. Apte, C. La Vecchia, C. D. Johnson, A. V. Biankin, et al., Pancreatic cancer. Nat. Rev. Dis. Prim. 2, 1–23 (2016).

2. L. Rahib, B. D. Smith, R. Aizenberg, A. B. Rosenzweig, J. M. Fleshman, L. M. Matrisian, Projecting cancer incidence and deaths to 2030: The unexpected burden of thyroid, liver, and pancreas cancers in the united states. Cancer Res. 74, 2913–2921 (2014).

3. N. C. Institute, Surveillance, Epidemiology, and End Results (SEER) Program Populations (1969–2015). DCCPS, Surveill. Res. Progr. (2017), (available at https://seer.cancer.gov).

4. M. A. Jacobetz, D. S. Chan, A. Neesse, T. E. Bapiro, N. Cook, K. K. Frese, et al., Hyaluronan impairs vascular function and drug delivery in a mouse model of pancreatic cancer. Gut. 62, 112–120 (2013).

5. K. P. Olive, M. a Jacobetz, C. J. Davidson, A. Gopinathan, D. McIntyre, D. Honess, et al., Inhibition of Hedgehog Signaling Enhances Delivery of Chemotherapy in a Mouse Model of Pancreatic Cancer. Science (80-. ). 324, 1457–1461 (2009).

6. P. P. Provenzano, C. Cuevas, A. E. Chang, V. K. Goel, D. D. Von Hoff, S. R. Hingorani, Enzymatic Targeting of the Stroma Ablates Physical Barriers to Treatment of Pancreatic Ductal Adenocarcinoma. Cancer Cell. 21, 418–429 (2012).

7. B. C. Özdemir, T. Pentcheva-Hoang, J. L. Carstens, X. Zheng, C. C. Wu, T. R. Simpson, et al., Depletion of carcinoma-associated fibroblasts and fibrosis induces immunosuppression and accelerates pancreas cancer with reduced survival. Cancer Cell. 25, 719–734 (2014).

8. A. D. Rhim, P. E. Oberstein, D. H. Thomas, E. T. Mirek, C. F. Palermo, S. A. Sastra, et al., Stromal elements act to restrain, rather than support, pancreatic ductal adenocarcinoma. Cancer Cell. 25, 735–747 (2014).

9. H. xiang Zhan, B. Zhou, Y. gang Cheng, J. wei Xu, L. Wang, G. yong Zhang, et al., Crosstalk between stromal cells and cancer cells in pancreatic cancer: New insights into stromal biology. Cancer Lett. 392, 83–93 (2017).

10. D. Von Ahrens, T. D. Bhagat, D. Nagrath, A. Maitra, A. Verma, The role of stromal cancer-associated fibroblasts in pancreatic cancer. J. Hematol. Oncol. 10, 1–8 (2017).

11. C. Liang, S. Shi, Q. Meng, D. Liang, S. Ji, B. Zhang, et al., Complex roles of the stroma in the intrinsic resistance to gemcitabine in pancreatic cancer: Where we are and where we are going. Exp. Mol. Med. 49 (2017), doi:10.1038/emm.2017.255.

12. C. J. Tape, S. Ling, M. Dimitriadi, K. M. McMahon, J. D. Worboys, H. S. Leong, et al., Oncogenic KRAS Regulates Tumor Cell Signaling via Stromal Reciprocation. Cell. 165, 910–920 (2016).

13. K. E. Richards, A. E. Zeleniak, M. L. Fishel, J. Wu, L. E. Littlepage, R. Hill, Cancer-associated fibroblast exosomes regulate survival and proliferation of pancreatic cancer cells. Oncogene. 36, 1770–1778 (2017).

14. T. S. Mantoni, S. Lunardi, O. Al-Assar, A. Masamune, T. B. Brunner, Pancreatic stellate cells radioprotect pancreatic cancer cells through β1-integrin signaling. Cancer Res. 71, 3453–3458 (2011).

15. H. Zhang, H. Wu, J. Guan, L. Wang, X. Ren, X. Shi, et al., Paracrine SDF-1&alpha; signaling mediates the effects of PSCs on GEM chemoresistance through an IL-6 autocrine loop in pancreatic cancer cells. Oncotarget. 6, 3085–3097 (2015).

16. L. Ireland, A. Santos, M. S. Ahmed, C. Rainer, S. R. Nielsen, V. Quaranta, et al., Chemoresistance in pancreatic cancer is driven by stroma-derived insulin-like growth factors. Cancer Res. 76, 6851–6863 (2016).

17. H. Miyamoto, T. Murakami, K. Tsuchida, H. Sugino, H. Miyake, S. Tashiro, Tumor-stroma interaction of human pancreatic cancer: acquired resistance to anticancer drugs and proliferation regulation is dependent on extracellular matrix proteins. Pancreas. 28, 38–44 (2004).

18. A. Neesse, C. Verbeke, E. Hessmann, L. Klein, F. M. Richards, A. Gopinathan, et al., Fibroblast drug scavenging increases intratumoural gemcitabine accumulation in murine pancreas cancer. Gut. 67, 497–507 (2017).

19. W. Plunkett, P. Huang, C. E. Searcy, V. Gandhi, Gemcitabine: preclinical pharmacology and mechanisms of action. Semin. Oncol. 23, 3–15 (1996).

20. J. D. Young, S. Y. M. Yao, J. M. Baldwin, C. E. Cass, S. A. Baldwin, The human concentrative and equilibrative nucleoside transporter families, SLC28 and SLC29. Mol. Aspects Med. 34, 529–547 (2013).

21. E. Mini, S. Nobili, B. Caciagli, I. Landini, T. Mazzei, Cellular pharmacology of gemcitabine. Ann. Oncol. 17, 7–12 (2006).

22. E. A. Collisson, A. Sadanandam, P. Olson, W. J. Gibb, M. Truitt, S. Gu, et al., Subtypes of pancreatic ductal adenocarcinoma and their differing responses to therapy. Nat. Med. 17, 500–503 (2011).

23. R. F. Knoop, M. Sparn, J. Waldmann, L. Plassmeier, D. K. Bartsch, M. Lauth, et al., Chronic Pancreatitis and Systemic Inflammatory Response Syndrome Prevent Impact of Chemotherapy with Gemcitabine in a Genetically Engineered Mouse Model of Pancreatic Cancer. Neoplasia. 16, 463–470 (2014).

24. M. G. Bachem, E. Schneider, H. Groß, H. Weidenbach, R. M. Schmid, A. Menke, et al., Identification, culture, and characterization of pancreatic stellate cells in rats and humans. Gastroenterology. 115, 421–432 (1998).

25. P. Mews, P. Phillips, R. Fahmy, M. Korsten, R. Pirola, J. Wilson, et al., Pancreatic stellate cells respond to inflammatory cytokines: potential role in chronic pancreatitis. Gut. 50, 535–541 (2002).

26. S. Y. Lunt, V. Muralidhar, A. M. Hosios, W. J. Israelsen, D. Y. Gui, L. Newhouse, et al., Article Pyruvate Kinase Isoform Expression Alters Nucleotide Synthesis to Impact Cell Proliferation, 95–107 (2015).

27. M. Hafner, M. Niepel, M. Chung, P. K. Sorger, Growth rate inhibition metrics correct for confounders in measuring sensitivity to cancer drugs. Nat. Methods. 13, 1–11 (2016).

28. J. Ciccolini, C. Serdjebi, G. J. Peters, E. Giovannetti, Pharmacokinetics and pharmacogenetics of Gemcitabine as a mainstay in adult and pediatric oncology: an EORTC-PAMM perspective. Cancer Chemother. Pharmacol. 78, 1–12 (2016).

29. M. Klanova, L. Lorkova, O. Vit, B. Maswabi, J. Molinsky, J. Pospisilova, et al., Downregulation of deoxycytidine kinase in cytarabine-resistant mantle cell lymphoma cells confers cross-resistance to nucleoside analogs gemcitabine, fludarabine and cladribine, but not to other classes of anti-lymphoma agents. Mol. Cancer. 13, 1–14 (2014).

30. T. Qin, J. Jelinek, J. Si, J. Shu, J. P. J. Issa, Mechanisms of resistance to 5-aza-2’-deoxycytidine in human cancer cell lines. Blood. 113, 659–667 (2009).

31. B. Nicander, P. Reichard, Evidence for the involvement of substrate cycles in the regulation of deoxyribonucleoside triphosphate pools in 3T6 cells. J. Biol. Chem. 260, 9216–9222 (1985).

32. V. Bianchi, E. Pontis, P. Reichard, Interrelations between substrate cycles and de novo synthesis of pyrimidine deoxyribonucleoside triphosphates in 3T6 cells. Proc. Natl. Acad. Sci. U. S. A. 83, 986–90 (1986).

33. V. Bianchi, E. Pontis, P. Reichard, Regulation of pyrimidine deoxyribonucleotide metabolism by substrate cycles in dCMP deaminase-deficient V79 hamster cells. Mol Cell Biol. 7, 4218–4224 (1987).

34. P. M. Bruno, Y. Liu, G. Y. Park, J. Murai, C. E. Koch, T. J. Eisen, et al., A subset of platinum-containing chemotherapeutic agents kills cells by inducing ribosome biogenesis stress. Nat. Med. 23, 461–471 (2017).

35. A. L. Mccleary-Wheeler, R. Mcwilliams, M. E. Fernandez-Zapico, Aberrant signaling pathways in pancreatic cancer: A two compartment view. Mol. Carcinog. 51, 25–39 (2012).

36. N. A. Ottenhof, R. F. de Wilde, A. Maitra, R. H. Hruban, G. J. A. Offerhaus, Molecular Characteristics of Pancreatic Ductal Adenocarcinoma. Patholog. Res. Int. 2011, 1–16 (2011).

37. D. P. Ryan, T. S. Hong, N. Bardeesy, Pancreatic Adenocarcinoma. N. Engl. J. Med. 371, 1039–1049 (2014).

